# Temperature Influences Commensal-Pathogen Dynamics in a Nasal Epithelial Cell Co-culture Model

**DOI:** 10.1101/2023.10.06.561218

**Authors:** Joshua T. Huffines, RaNashia L. Boone, Megan R. Kiedrowski

## Abstract

Chronic rhinosinusitis (CRS) is an inflammatory disease of the paranasal sinuses, and microbial dysbiosis associated with CRS is thought to be a key driver of host inflammation that contributes to disease progression. *Staphylococcus aureus* is a common upper respiratory tract (URT) pathobiont that is associated with higher carriage rates in CRS populations, where *S. aureus* secreted toxins can be identified in CRS tissue samples. Although many genera of bacteria colonize the URT, relatively few account for the majority of sequencing reads. These include *S. aureus*, as well as several species belonging to the genus *Corynebacterium*, including *Corynebacterium propinquum* and *Corynebacterium pseudodiphtheriticum*, which are observed at high relative abundance in the URT of healthy individuals. Studies have examined the bacterial interactions between the major microbionts of the URT and *S. aureus*, but few have done so in the context of a healthy versus diseased URT environment. Here, we examine the role of temperature in commensal, pathogen, and epithelial dynamics using an air-liquid interface cell culture model mimicking the nasal epithelial environment. The healthy URT temperature changes from the nares to the nasopharynx and is altered during disease. Temperatures representative of the healthy URT increase persistence and aggregate formation of commensal *C. propinquum and C. pseudodiphtheriticum*, reduce *S. aureus* growth, and lower epithelial cytotoxicity compared to higher temperatures correlating with the diseased CRS sinus. Dual-species colonization revealed species-specific interactions between commensal *Corynebacterium* species and *S. aureus* dependent on temperature. Our findings suggest that URT mucosal temperature plays a significant role in mediating polymicrobial and host-bacterial interactions that may exacerbate microbial dysbiosis found in chronic URT disease.

**IMPORTANCE:** Chronic rhinosinusitis is a complex inflammatory disease with a significant healthcare burden. Although presence of *S. aureus* and microbial dysbiosis are considered mediators of inflammation in CRS, no studies have examined the influence of temperature on *S. aureus* interactions with the nasal epithelium and the dominant genus of the healthy URT, *Corynebacterium*. Interactions between *Corynebacterium species* and *S. aureus* have been documented in several studies, but none to date have examined how environmental changes in the URT may alter their interactions with the epithelium or each other. This study utilizes a polarized epithelial cell culture model at air-liquid interface to study the colonization and spatial dynamics of *S. aureus* and clinical isolates of *Corynebacterium* from people with CRS to characterize the role temperature has in single-and dual-species dynamics on the nasal epithelium.

## INTRODUCTION

Chronic rhinosinusitis (CRS) is a highly prevalent and costly inflammatory disease of the upper respiratory tract (URT) with up to 12% of the United States population affected, leading to annual costs approaching $19 billion (1–3). Inflammation is central to CRS pathology, and microbial dysbiosis in the sinuses and URT is considered a key contributor to chronic inflammation, furthering disease progression (4–6). The healthy URT microbiome is highly diverse, but most sequencing reads come from relatively few bacterial genera (7). Reduced microbial diversity is a hallmark of many airway diseases, yet for CRS findings have been inconclusive, with some studies reporting reduced diversity in CRS populations and others increased diversity (4). One perspective is that CRS dysbiosis is not necessarily driven by a reduction in diversity, but either a shift from commensal microbes to pathogenic ones or instigation of pathogenic behavior from pathobionts. Among the prevalent and abundant members of the URT microbiome, *Staphylococcus aureus* is commonly associated with CRS and has higher carriage rates in CRS patient populations (8–10). CRS inflammatory responses have been linked to *S. aureus*-specific toxins, and *S. aureus* toxins and antibodies to *S. aureus* secreted factors can be identified in CRS tissue samples (11–14). Longitudinal carriage of *S. aureus* is associated with worsening CRS symptoms, and carriage is also linked to disease recurrence after endoscopic sinus surgery (15, 16). Interestingly, antagonistic interactions with *S. aureus* have been documented for several prevalent and abundant members of the healthy URT microbiome (17–24). Despite this, *S. aureus* can reach an average relative abundance nearing 48% of the sinonasal microbiome in the presence of these competing microbionts (25). No studies to date have investigated the environmental factors leading to *S. aureus* dominance over other microbionts within the context of CRS.

Differences in nasal mucosal temperature may influence the commensal, pathogen, and host dynamics in CRS. The nasal mucosa of healthy people has been reported to vary between 29°C and 32°C for the nares and nasal turbinates (26, 27). However, the nasal cavity undergoes significant changes with the onset of CRS, including significant mucus production, mucosal swelling, possible polyp formation, and decreased epithelial barrier integrity (28). Consequently, nasal obstruction and decreased airflow are often seen in CRS patients, which can be alleviated through endoscopic sinus surgery (3, 29–32). Additionally, inflammation and bacterial infection, key components of CRS pathology, have been shown to increase skin temperatures in post-surgery wounds and skin ulcers (33, 34). Reduced airflow and inflammation are both thought to increase the temperature of the nasal mucosa (35). Supporting this theory, thermographic imaging of CRS patients has successfully been used as a strategy to identify which nasal passage or sinus cavity is affected due to significantly increased temperatures (36–38). This difference in temperature can have profound effects on bacterial growth and behavior. Indeed, *S. aureus* has been shown to induce significant changes in its transcriptome and proteome when grown in rich-medium at temperatures ranging from 34-40°C, with higher temperatures increasing its hemolytic capability (39).

In this study, we examine how temperature affects the interactions between nasal epithelial cells, *S. aureus,* and two CRS clinical isolates of *Corynebacterium*, the most abundant genus in the healthy URT (7), using an air-liquid interface model of human nasal epithelial cells. *Corynebacterium* species are significantly depleted in CRS, and several studies have found *Corynebacterium* is negatively correlated with *S. aureus* on skin and in the nares (7, 25, 40–42). Additionally, some *Corynebacterium* species have been shown to reduce *S. aureus* virulence and potentially induce autolysis, which has made *Corynebacterium* a candidate for probiotic usage (17, 23, 43–46). Understanding how environmental changes affect the microbial constituents of the URT is important for elucidating the changes in the microbiome observed in disease, including the growth of *S. aureus*. We hypothesize that increasing temperature leads to a favorable environment for *S. aureus* to outcompete *Corynebacterium* on the URT epithelium. This study characterizes the role of temperature in commensal, pathogen, and nasal epithelium dynamics, shedding light on the complex dynamics of chronic URT disease pathology.

## RESULTS

### Lower temperature improves Corynebacterium pseudodiphtheriticum and Corynebacterium propinquum persistence but limits S. aureus growth in culture

To assess the impact of temperature on *S. aureus* and *Corynebacterium*, we first evaluated bacterial growth in rich liquid culture medium. For *S. aureus,* we evaluated the thoroughly studied methicillin-resistant strain USA300 LAC that is representative of difficult-to-treat *S. aureus* strains encountered in CRS (47). We utilized two clinical isolates of *Corynebacterium* from CRS patients (16, 48), *Corynebacterium propinquum* and *Corynebacterium pseudodiphtheriticum*, due to their prevalence and high relative abundance in the URT (7). We also tested the *Corynebacterium glutamicum* strain ATCC 13032, a well-characterized and commonly used industrial strain, to serve as a comparison for the airway-adapted *Corynebacterium* clinical isolates’ growth *in vitro*. Each of these strains was grown in 96-well microtiter plate with brain-heart infusion (BHI) broth for up to 48 hours at 37°C and 30°C (Figure 1). Absorbance at 600nm was measured at 4, 8, 12, 24, and 48 hours, and bacteria were harvested to determine viable colony-forming units (CFUs) at each timepoint. *S. aureus* absorbance values were significantly higher at 37°C at each timepoint measured after the initial reading, and CFU counts showed a modest reduction in CFUs at lower temperature (Figure 1A). To test whether this trend is seen in other strains of *S. aureus*, including isolates from CRS patients, we examined the growth differences of USA100 and three CRS isolates of *S. aureus* at 37°C and 30°C (Supplementary Figure 2) which matched the trends seen in Figure 1A. *C. glutamicum* CFUs were significantly lower at 30°C early during growth but overall, both measurements for *C. glutamicum* showed little difference between growth at low and high temperatures over time (Figure 1B). Unlike *C. glutamicum*, *C. propinquum* and *C. pseudodiphtheriticum* CRS isolates had significantly greater CFUs at 24-and 48-hour timepoints when grown at 30°C and significantly increased absorbance readings at 30°C throughout the assay, with the greatest differences observed at later timepoints (Figure 1CD).

**Figure 1:**
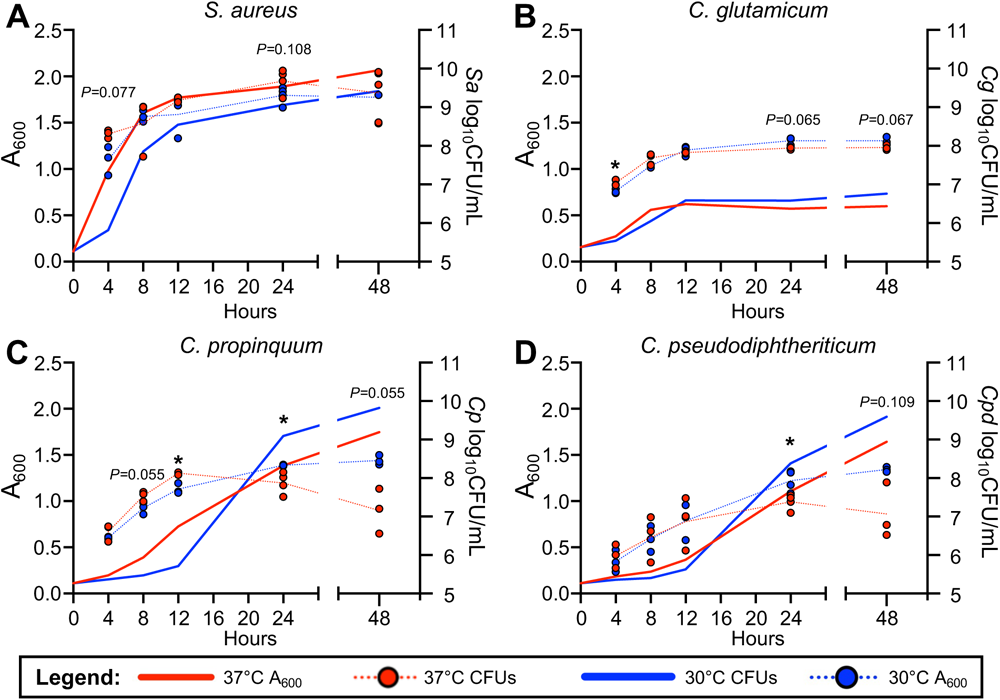
Lower temperatures increase persistence of clinical *Corynebacterium* isolates. Absorbance readings and viable colony-forming units for *S. aureus* USA300 LAC (A), *C. glutamicum* ATCC 13032 (B), *C. propinquum* (C), and *C. pseudodiphtheriticum* (D) grown in brain-heart infusion broth at either 37°C (red) or 30°C (blue) for 48 hours.

### Lower temperatures boost *C. propinquum and C. pseudodiphtheriticum* persistence on HNECs while limiting *S. aureus* growth

To examine whether trends observed in vitro in liquid culture also apply to bacterial growth in association with the nasal epithelium, we tested adherence and growth of *S. aureus*, *C. propinquum* and *C. pseudodiphtheriticum* on polarized human nasal epithelial cells (HNECs) at air-liquid interface at 37°C and 30°C (Figure 2). *S. aureus* adherence at 1 hour was unaffected by temperature, however growth on HNECs after 6 hours was significantly lower at 30°C compared to the 37°C group (Figure 2A). Additionally, we measured growth on HNECs for USA100 and three CRS isolates of *S. aureus* (Supplementary Figure 3) which matched the trend seen here. Temperature did not greatly affect *C. propinquum and C. pseudodiphtheriticum* adherence or growth at 6 hours (Figure 2BC). However, at later timepoints evaluated *C. propinquum* had significantly higher CFUs at 24-hours at 30°C (Figure 2B), and *C. pseudodiphtheriticum* had a similar increase in growth at 30°C at both the 18-and 24-hour timepoints (Figure 2C).

**Figure 2:**
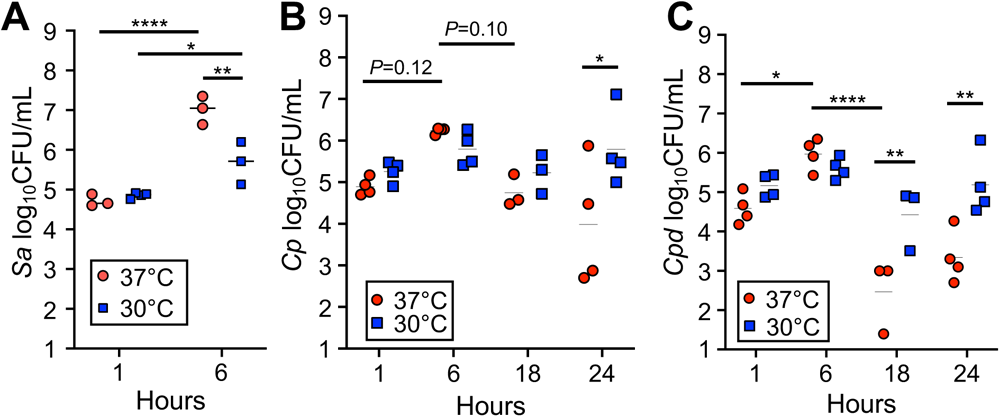
Lower temperatures support *Corynebacterium* persistence but reduce *S. aureus* growth on human nasal epithelial cells (HNECs) at air-liquid interface. *S. aureus* (A) viable colony-forming units from growth on polarized RPMI2650 HNECs at air-liquid interface for 1 and 6 hours at 37°C (red) or 30°C (blue). *C. propinquum* (B) and *C. pseudodiphtheriticum* (C) viable colony-forming units from grown on polarized RPMI2650 nasal epithelial cells at 37°C (red) or 30°C (blue) for 1, 6, 18, and 24 hours. n=3-6 biological replicates. Two-way ANOVA *(*P<*=0.05,*\*\*P<*=0.01, \*\*\**P*<=0.001, \*\*\*\**P*<=0.0001).

Interestingly, we observed both of the *Corynebacterium* clinical isolates had lower CFUs at 18 hours compared to 6 hours when grown at the higher 37°C temperature (Figure 2BC). To ascertain whether these differences stemmed from altered aggregation on the surface of HNEC cultures, we utilized fluorescence microscopy to visualize *Corynebacterium* growth on HNECs. We modified the pJOE7706.1 plasmid backbone (49) to express the red fluorescent protein, tdTomato, under the control of an IPTG-inducible promoter. We then modified a transformation protocol developed for *C. glutamicum* and electroporated pJOE7706.1-tdTomato into *C. propinquum* and *C. pseudodiphtheriticum* to obtain fluorescent strains for microscopy. We inoculated HNECs with each isolate and grew them at 37°C and 30°C for 6, 18, and 24 hours, omitting the 1-hour timepoint due to negligible differences in CFUs. *Corynebacterium* aggregate formation on HNECs appeared substantially increased at 30°C at the 18-and 24-hour timepoints (Figure 3AB). Consistent with the HNEC CFU experiments (Figure 2BC), both *Corynebacterium* isolates had reduced aggregate size at later timepoints observed for the 37°C group (Figure 3AB). 3D volume views of z-stacks acquired for the 24-hour timepoint showed both *C. propinquum* and *C. pseudodiphtheriticum* isolates grew on the apical surface of the epithelial layer (Figure 3CD). To assess whether *S. aureus* followed the same trends observed via CFU counts, we repeated colonization of HNECs at the 6-hour timepoint with GFP-expressing *S. aureus* (50). Coverage of the HNEC surface was largely reduced at 30°C (Figure 3E) consistent with CFU data. Notably, while *S. aureus* aggregate formation and size appeared to be mildly reduced at 30°C, the majority of reduction in *S. aureus* appeared to be from an almost complete lack of bacteria not in large clusters or aggregates (Figure 3E). Similar to *Corynebacterium*, *S. aureus* aggregates were found on the surface of the epithelial layer (Figure 3E).

**Figure 3:**
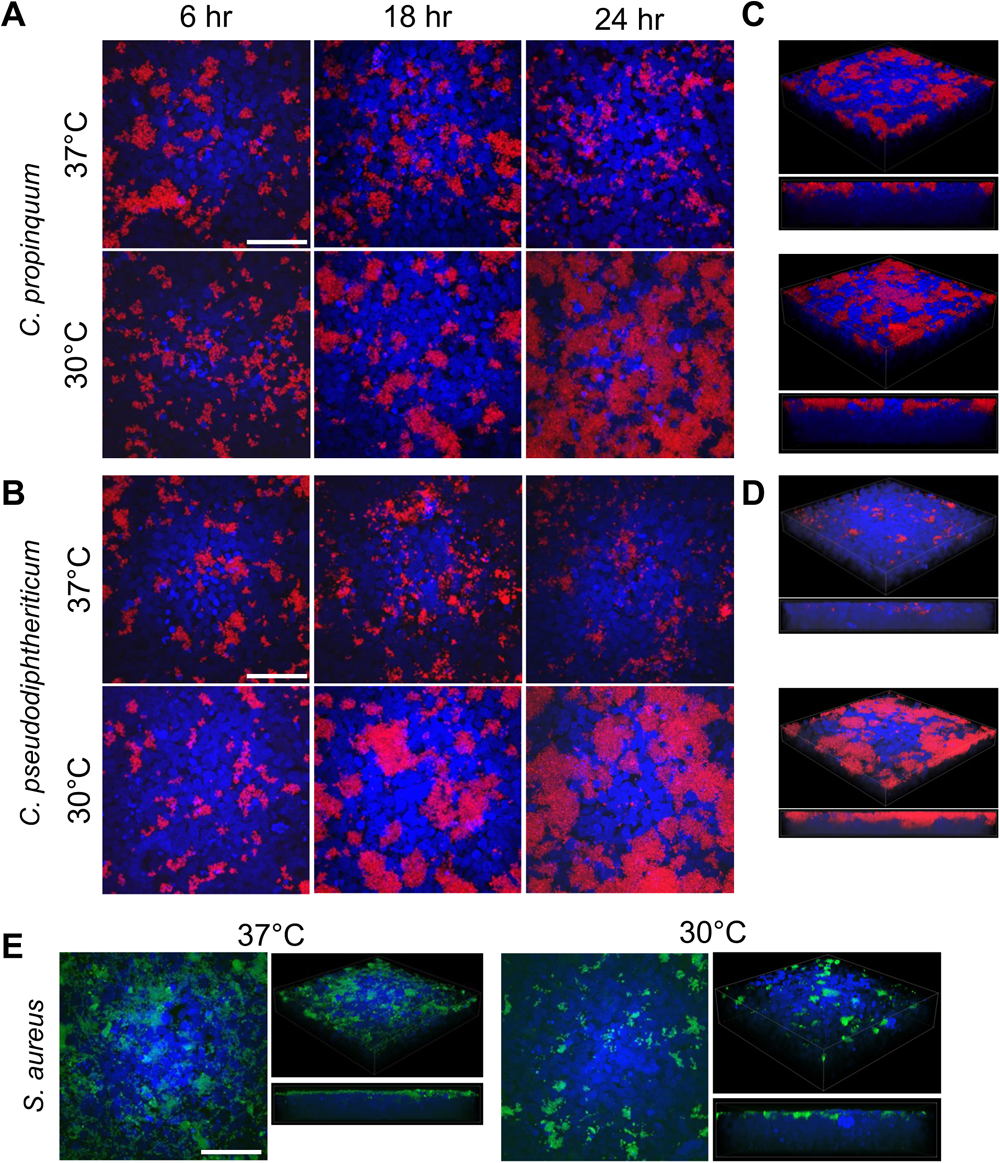
Lower temperatures support large aggregate formation of *Corynebacterium* on human nasal epithelial cells (HNECs) and reduce *S. aureus* colonization. 2D Fluorescence microscopy of tdTomato-expressing *C. propinquum* (A) and *C. pseudodiphtheriticum* (B) on polarized air-liquid interface HNECs stained with Hoechst grown at 37°C or 30°C for 6, 18, and 24 hours. 3D and side views of the 24-hour images for *C. propinquum* (C) and *C. pseudodiphtheriticum* (D) on Hoechst-stained HNECs. 2D and 3D fluorescence microscopy images of GFP-expressing *S. aureus* (E) grown on HNECs for 6 hours.

### Temperature alters polymicrobial growth on HNECs

Considering how low and high temperatures affected growth of each species alone, we next asked how temperature may modulate interactions between *S. aureus* and *Corynebacterium* when grown in co-culture on HNECs, an environment that more closely represents the polymicrobial setting in the URT. We developed two models to investigate co-inoculation and sequential inoculation with *S. aureus* and each *Corynebacterium* CRS isolate (Supplementary Figure 1) and characterized the effects of incubation at 37°C or 30°C on dual-species colonization of HNECs. *S. aureus* CFUs remained significantly decreased at 30°C when grown with either *C. propinquum or C. pseudodiphtheriticum* using the co-inoculation scheme (Figure 4A). *S. aureus* growth in co-culture did not differ substantially from *S. aureus* grown alone on HNECs, as determined by measuring the fold-change in CFUs from dual-species culture normalized to CFUs from single-species culture on HNECs (Figure 4C). Sequential inoculation did not result in significant changes in *S. aureus* CFUs at either temperature (Figure 4A). In contrast to *S. aureus,* growth of both *Corynebacterium* isolates benefitted from sequential inoculation at 37°C, with *C. propinquum* having the largest increase from single-species culture (Figure 4C). Here, no trend was observed due to temperature for *Corynebacterium* species and temperature (Figure 4BC).

**Figure 4:**
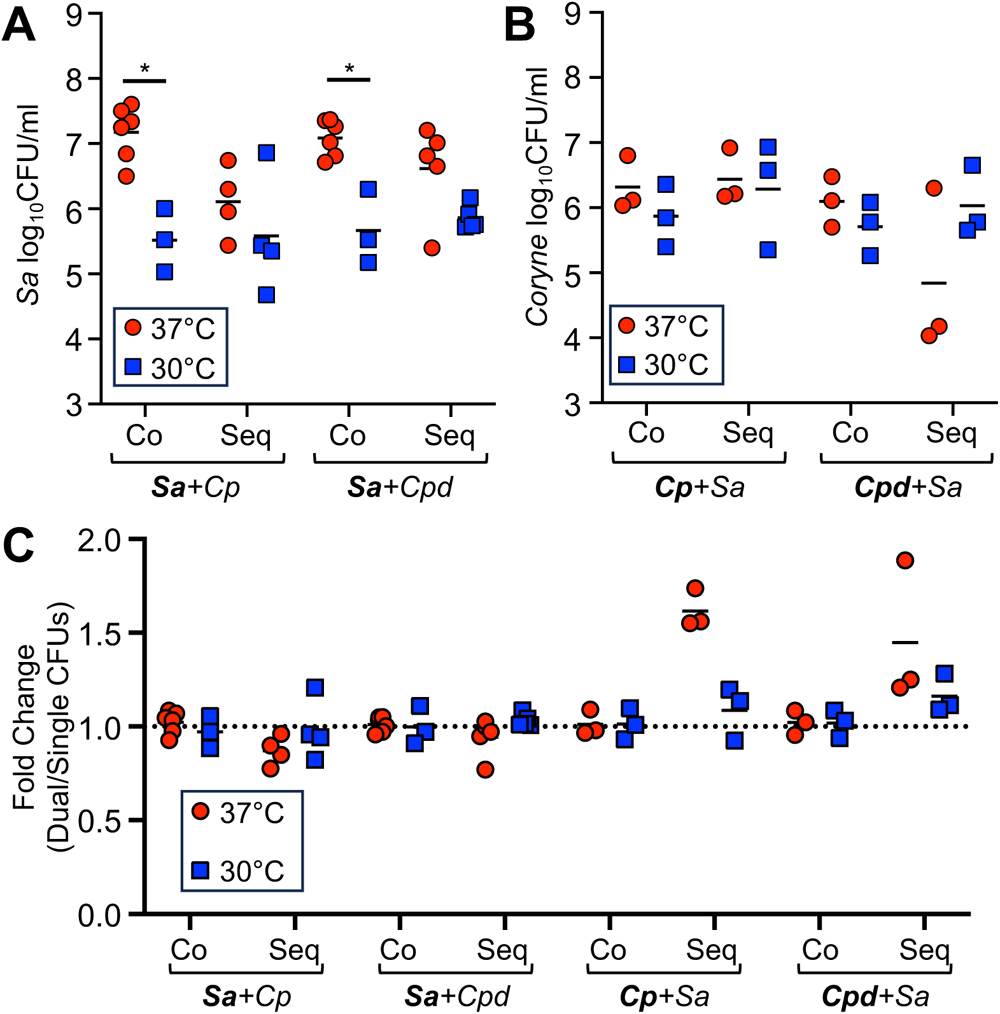
Dual-species bacterial colonization of human nasal epithelial cells demonstrates species-specific interactions with temperature. *S. aureus* (A) viable colony-forming units on HNECs from 6-hour co-inoculation or 6-hour sequential inoculation. *Corynebacterium* (B) viable colony-forming units on HNECs from 6-hour co-inoculation or 24-hour sequential inoculation with *S. aureus.* Fold-changes (C) for the dual-species CFUs normalized to the averages from single-culture CFUs.

We next examined growth following dual-species sequential inoculation using fluorescence microscopy (Figure 5). Strikingly, sequential inoculation with *S. aureus* improved growth of *C. propinquum* on HNECs at 37°C at the 24-hour timepoint (Figure 5A) supporting the increase in CFUs seen in Figure 4. In contrast, *C. pseudodiphtheriticum* grew similarly in the presence of *S. aureus* as observed in single species assays (Figure 3B), although with aggregates appeared smaller at 30°C in sequential inoculation with *S. aureus* than when grown alone (Figure 5B). *S. aureus* growth on HNECs appeared substantially reduced when grown with *C. propinquum* at either temperature and with *C. pseudodiphtheriticum* at 30°C (Figure 5).

**Figure 5:**
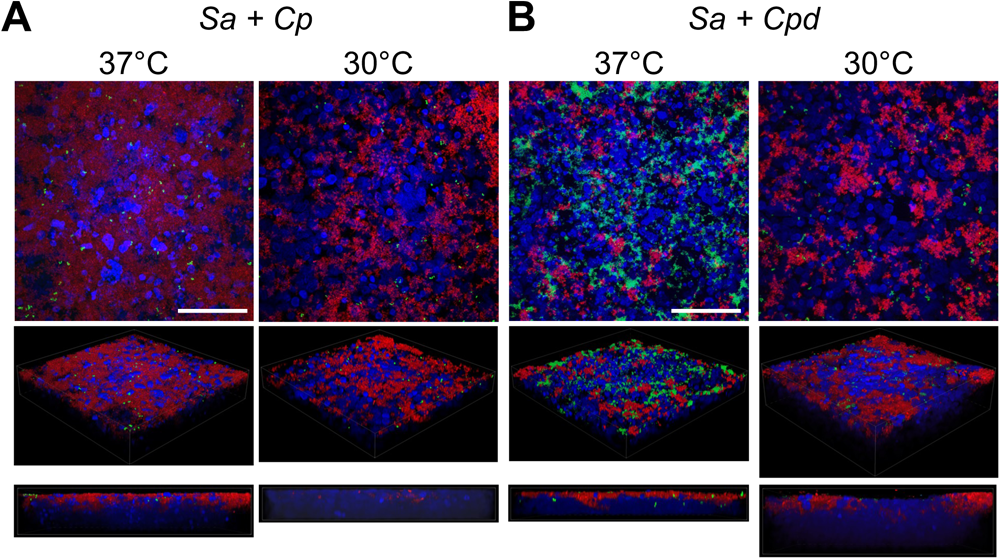
*Corynebacterium* has species-specific interactions with *S. aureus* at higher temperatures. Dual-species culture on human nasal epithelial cells (HNECs) at air-liquid interface with GFP-expressing (green) *S. aureus* and tdTomato-expressing (red) *C. propinquum* (A) or *C. pseudodiphtheriticum* (B) grown using the sequential model of inoculation at either 37°C or 30°C.

### Bacterial-induced cytotoxicity towards HNECs is reduced at lower temperatures

Having established single-and dual-species assays to evaluate growth of *Corynebacterium* and *S. aureus* on HNECs, we next examined the role temperature may have in modulating cytotoxicity of each species towards the epithelium. Basolateral media was collected from single-and dual-species colonized HNECs from sequential model of inoculation with fresh cell culture medium added 6 hours prior to harvest. Using a lactate dehydrogenase release assay to test the basolateral media, we determined epithelial cytotoxicity for uninfected controls and colonized groups. Surprisingly, *S. aureus* cytotoxicity was not significantly altered between temperatures (Figure 6) despite the significantly lower CFUs observed at 30°C (Figure 2A). *C. propinquum* showed significantly increased cytotoxicity at 37°C compared to 30°C, although colonization was increased at lower temperatures as observed through CFU counts. Using a mixed-model analysis to examine interactions between species groups and temperature, temperature was found to significantly reduce cytotoxicity (*P*=0.001) independently of the bacterial species colonizing the HNECs (Figure 6).

**Figure 6:**
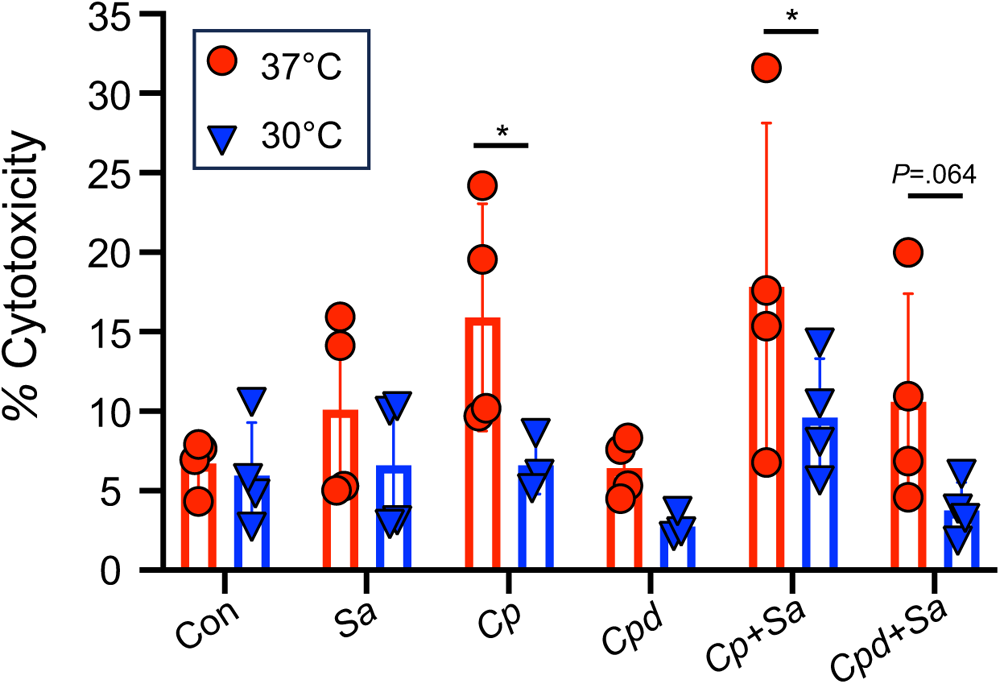
Lower temperatures reduce human nasal epithelial cell (HNEC) cytotoxicity independent of *S. aureus* and *Corynebacterium*. Lactate dehydrogenase release assay on the basolateral medium of HNECs colonized with *S. aureus, C. propinquum, C. pseudodiphtheriticum*, or dual-species using the sequential model of inoculation. HNECs were incubated with bacteria at either 37°C (red) or 30°C (blue).

## DISCUSSION

In this study, we sought to characterize the role temperature may play in modulating interactions between bacterial constituents of the URT in the presence of the nasal epithelium. The temperature of the healthy nasal epithelium is approximately 30°C, and this temperature increases during chronic inflammatory URT disease (26, 27, 35, 37, 51, 52). Given the microbial dysbiosis commonly observed in CRS populations (53), we hypothesized that increased temperatures in the diseased URT may benefit one of the major CRS pathogens, *S. aureus* (9, 12–14, 16), allowing for it to outcompete commensal bacteria. Here, we evaluated how temperature affects a well-studied MRSA strain alongside two species of *Corynebacterium* collected from people with CRS (16) using a human epithelial cell co-culture model.

By first testing single-species of growth in rich culture medium and on HNECs, we found the clinical isolates of *Corynebacterium* had reduced persistence at a higher temperature of 37°C compared to 30°C (Figs. 1CD and 2BC). In contrast, *S. aureus* growth benefited at a higher temperature, which supported our initial hypothesis. Fluorescence microscopy supported these findings, showing substantial increases in *Corynebacterium* growth and reduction in *S. aureus* growth on nasal epithelial cells incubated at the lower temperature of 30°C (Fig. 3). Considering these findings in the context of colonization and pathogenicity, *S. aureus* is a notorious pathogen known to infect numerous body sites where the temperature is at or above 37°C, leading to osteomyelitis, sepsis, soft tissue infections and pneumonia (10). *S. aureus* transcriptional changes between 34°C and 40°C have been investigated (39) in rich medium, and significant increases in hemolytic activity at higher temperatures were observed, indicative of increased pathogenic potential at higher temperatures. Here, our study investigated the effects of temperature on *S. aureus* behavior when in the presence of epithelial cells, with the nutritional environment originating solely from epithelial sources, at temperatures representative of the healthy nasal mucosa and the impacted sinus in CRS disease. Although increased protease expression has been observed at lower temperatures in BHI (39), and protease activity is implicated in biofilm dispersal (54), *S. aureus* was mostly found in large aggregates on the surface of epithelial cells at lower temperatures (Fig. 3E). However, the nature of *S. aureus* biofilm formation may vary in different environments, relying on other secreted factors for dispersal such as nuclease and phenol-soluble modulins (PSMs) (55, 56), with PSMs known to be more highly expressed at higher temperatures (39).

For *Corynebacterium*, the effects of temperature on growth and metabolism of *C. glutamicum* have been previously studied to optimize growth of this environmental strain used in industry for production of amino acids, and lower temperatures were found to benefit its growth over longer time periods (57–59). Our data supports this observation for CRS clinical isolates of *Corynebacterium*, with growth and persistence of *C. propinquum* and *C. pseudodiphtheriticum* both found to be higher at the lower temperature of 30°C compared to 37°C in rich medium and on epithelial cells (Figs. 1CD and 2BC). No studies to our knowledge have investigated the effect of lower temperatures representative of the healthy URT mucosal surface on commensal URT *Corynebacterium* species and human epithelial cells. Additionally, although research on *Corynebacterium diphtheriae* identified adhesins important for colonizing the epithelium (60), there has been no characterization of biofilm or aggregate formation of this species or other corynebacteria on nasal epithelial surfaces over time, as we report here. Some *Corynebacterium* species are known to form aggregates (61), possibly due to the hydrophobic nature of their outer mycolic acid membrane, but it is unknown whether this occurs for commensal *Corynebacterium* in the URT. Our results suggest that *Corynebacterium* can develop large biofilm-like structures in the nutritional milieu of the URT, which may impact interactions between *Corynebacterium* and other nasal microbiome constituents.

When examining dual-species colonization of HNECs, viable colony-forming units compared to single-species controls were not markedly different for *S. aureus,* whereas *C. propinquum* had a substantial increase at 37°C when cultured in the sequential model of inoculation with *S. aureus* (Fig. 4). In contrast, *C. pseudodiphtheriticum* largely maintained a temperature-sensitive trend in the presence of *S. aureus*, continuing to benefit at the lower temperature of 30°C. Fluorescence microscopy supported these data, confirming increased *C. propinquum* aggregates associated with HNECs at 37°C in the presence of *S. aureus* (Fig. 4). It is possible the increase of *C. propinquum* when grown together with *S. aureus* at higher temperatures is due to changes in available nutrients in the airway surface liquid due to metabolites produced either by *S. aureus* or from epithelial cells, which could result either from altered secretion due to the presence of staphylococcal proteins or increased *S. aureus*-induced cytotoxicity. Of note, previous studies have found metabolic interactions between URT microbionts, and many of these interspecies interactions have been documented showing synergism, which may explain the interactions seen for *C. propinquum* in the presence of *S. aureus* (62, 63).

After focusing on how URT colonizing bacteria responded to growth at different temperatures, we next asked how the nasal epithelium would adjust to these temperatures and associated changes in bacterial growth by examining cytotoxicity. Intriguingly, *C. propinquum* co-culture induced higher levels of cytotoxicity than *C. pseudodiphtheriticum*, with significantly higher measurements observed at 37°C (Fig. 6), the temperature at which colonization for *C. propinquum* was lowest according to both CFUs and microscopy (Figs. 2B and 3A). This suggests that temperature may play a significant role in the host response to bacterial colonization, as the uninfected controls showed no noticeable temperature-dependent difference in cytotoxicity. Temperature is likely a key factor affecting bacterial behavior in the URT, with increased temperature potentially acting as an environmental switch that induces a more pathogenic phenotype even in species that are normally considered to be commensals such as *Corynebacterium*. Overall, these data indicate a role for temperature in both colonization and bacterial behavior of *C. propinquum and C. pseudodiphtheriticum* that is species-specific, as both isolates had similar growth at lower temperatures yet significant differences in epithelial cytotoxicity. Considering this in the context of CRS, it is possible that the native microbiome constituents in the URT could have a vastly different impact on inflammation depending on the microenvironment of the mucosal epithelium. Elucidating factors involved in species-specific epithelial cytotoxicity of *Corynebacterium* species, using isolates collected from the healthy and diseased URT, is a topic that warrants further study.

The work presented here addresses gaps in knowledge regarding how *S. aureus* interacts with *Corynebacterium* when colonizing the nasal epithelium and the role of temperature in regulating interspecies interactions in the URT, yet there are limitations to this study. Notably, the clinical isolates of *Corynebacterium* we evaluated were from subjects with CRS, and these strains could have acquired adaptations to the chronically diseased URT due to selective pressures encountered in the CRS sinuses that lead them to interact differently with *S. aureus*. Future studies would benefit from investigating *Corynebacterium* species that may be more representative of the healthy URT. The HNEC ALI culture model used here provides a controlled environment for studying bacterial interactions with the epithelium and the nutritional environment in the URT, however this model lacks significant components of the innate and adaptive immune system, as well as other cell types present in the URT in addition to the epithelium. Considering how temperature affects inflammatory responses in *in vivo* models of CRS could uncover additional interactions that occur between *S. aureus* and *Corynebacterium* to influence disease progression or highlight potential avenues for more effective therapy to address chronic infections.

## MATERIALS AND METHODS

### Bacterial strains and growth conditions

Bacterial strains and plasmids used in this work are summarized in Table 1. *S. aureus* and *Corynebacterium* strains were cultured overnight at 37°C with shaking in brain-heart infusion broth (BHI; BD Biosciences) unless otherwise noted. Cultures were inoculated using colonies grown on BHI with 1.5% agar (BD Biosciences) at 37°C. *E. coli* cultures were grown in Luria broth (LB) shaking at 37°C. The *S. aureus* USA300 LAC 13c clone used for all experiments was a gift from Tammy Kielian (64). The *E. coli* strain GM2163 was a gift from Chuck Turnbough. *Corynebacterium propinquum* and *Corynebacterium pseudodiphtheriticum* strains were isolated from the sinuses of subjects with cystic fibrosis and chronic rhinosinusitis by culturing from sinonasal swabs collected during endoscopic sinus procedures that were clinically indicated for management of patients’ chronic sinus disease (University of Pittsburgh IRB REN16110185) (16). Species identification was confirmed by PCR and sequencing of the 16s rRNA gene using primers 63f (5’-CAG GCC TAA CAC ATG CAA GTC-3’) and 1387r (5’-GGG CGG AGT GTA CAA GGC-3’) (65). Plasmids in *S. aureus* were maintained in 25 µg/ml of chloramphenicol (Sigma). Plasmids in *Corynebacterium* strains were maintained in 50 µg/mL of kanamycin (ThermoScientific). Plasmids in *E. coli* strains were maintained in 100 µg/ml of ampicillin (Fisher BioReagents).

**Table 1.** Bacterial strains and plasmids.

| Strain or Plasmid | Description | Source or Reference |
| --- | --- | --- |
| **Strains** |  |  |
| *S. aureus* LAC 13c | USA300 CA-MRSA, erm<sup>S</sup> | (64) |
| *S. aureus* RN4220 | Restriction deficient cloning host | (70) |
| *S. aureus* MRSA CRS isolates | Sinus clinical isolates | (16) |
| *Corynebacterium propinquum* | Sinus clinical isolate | (16) |
| *Corynebacterium pseudodiphtheriticum* | Sinus clinical isolate | (16) |
| *E. coli* DH5 alpha | Cloning strain | Zymo Research |
| *E. coli* GM2163 | Dam-/dcm- cloning strain | (68) |
| **Plasmids** |  |  |
| pCM29 | *S. aureus* sarAP1 promoter driving the sGFP gene | (71) |
| pJOE7706.1 | *Corynebacterium glutamicum* shuttle vector with eGFP containing a N-terminal his-tag | (49) |
| pJOE7706.1-tdtomato | pJOE7706.1 with eGFP swapped for tdtomato at BsrGI and BamHI sites | This work |
| pMQ361 | pBBR1-based shuttle vector with PnptII-tdtomato | (72) |

### Recombinant DNA and genetic techniques

Plasmid DNA was prepared from *E. coli* DH5-alpha using the NEB Monarch Plasmid Purification miniprep kit and then electroporated into *S. aureus* strain RN4220 as previously described (66). DNA was then moved from RN4220 into *S. aureus* LAC 13c through transduction with bacteriophage alpha-80 as previously described (67). For transformation into *Corynebacterium* strains, plasmid DNA was prepared from *E. coli* strain GM2163 (dam-/dcm-) (68). Restriction enzymes, polymerases were and enzymes for DNA modification purchased from New England Biolabs (Beverly, MA) and used according to manufacturer’s instructions. Oligonucleotides were synthesized by Integrated DNA Technologies (Coralville, IA). Non-radioactive sequencing was performed at the University of Alabama at Birmingham Heflin Center for Genomic Sciences Core Sequencing Facility.

#### Plasmid construction

Plasmid pJOE7706.1 was a gift from Josef Altenbuchner (Addgene plasmid # 135075; http://n2t.net/addgene:135075; RRID:Addgene_135075). To generate plasmid pJOE7706.1-tdtomato, plasmid pMQ361 was digested with BsrGI and BamHI (NEB) to obtain the tdtomato fragment, purified using the NEB Monarch DNA Gel Extraction kit, and then ligated to pJOE7706.1 cut with the same enzymes.

#### Preparation of electrocompetent *Corynebacterium*

Methods for preparing electrocompetent *Corynebacterium* were based on protocols described by Eggeling and Bott in the Handbook of *Corynebacterium glutamicum* (69). Electrocompetent *Corynebacterium* was made by starting a 5 mL overnight culture in BHIS medium, composed of BHI + 9.1% sorbitol (Fisher BioReagents), and culturing overnight at 30°C with shaking 200rpm. Next, 100 mL of BHIS was inoculated with 2-5 mL of the overnight culture to achieve a starting OD_600_ of 0.2, followed by 3-5 hours of growth at 30°C with shaking. Once an OD_600_ of 2.0 was reached, bacteria were centrifuged at 3400 RCF at 4°C. Cells were washed 3x with chilled 1mM Tris-HCl and 10% glycerol (Fisherbrand). Bacteria were then resuspended in chilled 10% glycerol and aliquoted for storage in at −80°C.

#### Transformation of *Corynebacterium*

Electroporation of *Corynebacterium*was accomplished by thawing aliquots of electrocompetent *Corynebacterium* on ice for 15 minutes prior to incubation with 200 ng of plasmid prepared from *E. coli* strain GM2163 for 15 minutes on ice. The mixture was transferred to a 2 mm gap electroporation cuvette (Fisherbrand) and electroporated using a BioRad Gene Pulser set at 2.5kV, 25µF, and 200Ω. After electroporation, 1 mL of BHIS was added followed by a 6-minute incubation in a 46°C water bath. Bacteria were then incubated at 30°C with shaking for 1 hour, followed by plating 100µL of bacterial suspension on BHIS agar plates containing 50 µg/mL kanamycin.

### Cell lines and growth conditions

The human nasal epithelial cell (HNEC) line RPMI2650 (ATCC CCL-30) was maintained in complete Eagle’s Minimum Essential Medium (Corning) containing L-glutamine (Gibco), Plasmocin (InvivoGen), Penicillin-Streptomycin (Gibco) and 10% fetal bovine serum (US qualified, Gibco) at 37°C and 5% CO_2_ unless otherwise noted. Adherent cells were washed once with DPBS prior to trypsinization with 0.25% trypsin + EDTA (Corning). Cells were collected by centrifugation at 1400 RCF for 3 minutes at 4°C and resuspended in media. Transwell inserts (Greiner Bio-One) were coated with Vitrogen Plating Medium (VPM), consisting of MEM without glutamine or phenol red (Gibco), 10 µg/mL fibronectin (Corning), 100 µg/mL of bovine serum albumin (Life Technologies) and 30 µg/mL of PureCol bovine collagen (Advanced Biomatrix), added to the apical side of the insert and crosslinked under UV light for a minimum of 45 minutes. VPM was then removed, and 2.5×10^5^ cells were added to the apical side of the insert. Apical and basolateral media was replaced every other day until 7 days after seeding, when apical media was removed for transition to air-liquid interface (ALI). After maintaining at ALI for 7 additional days, both apical and basolateral sides of the insert were washed with DPBS, and media without Plasmocin or Penicillin-Streptomycin was added to the basolateral side only for infection assays.

### Co-culture HNEC Assays

Overnight bacterial cultures were centrifuged at 3000RCF and washed once with sterile phosphate-buffered saline (Fisher). Bacteria were resuspended in MEM without glutamine or phenol red (Gibco) at an OD_600_ of 0.5. For inoculation of HNECs, *S. aureus* was further diluted in MEM to a final OD_600_ of 0.02, and *Corynebacterium* isolates were diluted to a final OD_600_ of 0.2. Then, 50 µL of bacterial suspension was added to the apical surface of the ALI HNEC cultures for 1 hour with incubation at 37°C or 30°C with 5% CO_2_. After one hour, apical media was removed and co-cultures were further incubated until 6-, 18-, or 24-hours post-inoculation was reached. Co-cultures were then washed once apically with MEM to remove non-adherent bacteria. Basolateral media was removed and aliquoted for cytotoxicity assays. A solution of MEM with 0.1% Triton X-100 (BioRad) was added to the transwell insert, followed by 15 minutes of orbital shaking at 250 rpm. Cells were scraped from the filter, and the cell suspension was transferred into 950µL of MEM with 0.1% Triton X-100 for a total of 1 mL. Tubes were gently vortexed for 3 minutes, followed by serial dilution plating to determine CFUs. For sequential inoculation of HNECs, *Corynebacterium* strains were inoculated as described above and incubated for 18 hours, followed by *S. aureus* inoculation and incubation for an additional 6 hours. Differential plating to obtain *S. aureus* and *Corynebacterium* CFU counts was accomplished by plating dual-species groups on mannitol salt agar (Oxoid) to enumerate *S. aureus* and BHI agar plates with 100 µg/mL fosfomycin (TCI) to enumerate *Corynebacterium*.

### Confocal Imaging of coculture biofilms

Biofilm assays were performed as described above, with the addition of 0.1 mM IPTG to MEM used for inoculation, using bacterial strains expressing GFP (*S. aureus* with pCM29) or tdTomato (*Corynebacterium* with pJOE7706.1-tdTomato). After each time point, basolateral media was removed, and the transwell insert was washed with Dulbecco’s phosphate-buffered saline (DPBS) and fixed overnight at 4°C with cold 4% paraformaldehyde (PFA; Electron Microscopy Sciences) diluted in DPBS. After fixation, PFA was removed, and samples were washed with DPBS followed by permeabilization with DPBS with 0.1% Triton X-100 for 15 minutes at room temperature. Samples then washed with DPBS followed by staining with a Hoechst (Invitrogen) diluted in DPBS for 15 minutes while shaking at room temperature. Samples were washed with DPBS, and filter inserts were cut out and mounted onto slides with ProLong Gold antifade reagent (Invitrogen). Microscopy was performed on a Nikon A1R microscopy system with the Nikon Eclipse Ti confocal microscope using the Plan Apo VC 60X Oil DIC N2 lens.

### Cytotoxicity Measurements

Basolateral media was collected from co-culture assays and stored at −80°C. For positive controls, 10 µL of 10X Lysis solution from the CytoTox 96 Non-Radioactive Cytotoxicity Assay (Promega) was added to 90 µL of basolateral media and incubated on the apical side of ALI cultures for 45 minutes before collecting. Basolateral media samples were thawed on ice prior to using the CytoTox 96 assay kit according to the manufacturer’s instructions. Cytotoxicity was calculated as a percentage value using cell culture medium as the negative control and fully lysed cells as the positive control. Uninfected cells cytotoxicity is displayed on the graph as the experimental negative control.

### Statistical analyses

Statistical analyses were performed with Graph Pad Prism version 9 software (GraphPad by Dotmatics). Two-way ANOVA with pre-determined multiple comparisons were used to evaluate the statistical significance between temperature and either time or bacterial group. Non-parametric unpaired two-tailed t-tests were used to evaluate the significance between colony-forming units for growth curves in rich-medium. The *P* value of 0.05 was used as the cutoff for statistical significance and *P* values below 0.10 are indicated on the graph with the specific value shown.

## ACKNOWLEDGEMENTS

Funding for this work was provided by the Cystic Fibrosis Foundation (KIEDRO18F5 and ROWE21R3) and the NIH (5T32GM008111-35).

---

**Supplementary Figure 1: Co-inoculation and sequential model of inoculation on HNECs.** Co-inoculation (top) and sequential (bottom) inoculation models of dual-species colonization on polarized HNECs at air-liquid interface.

**Supplementary Figure 2: Lower temperatures reduce growth in *S. aureus* isolates from chronic rhinosinusitis patients.** Growth of *S. aureus* strains in brain-heart infusion broth at 37°C (red) and 30°C (blue) for 24 hours. 3 biological replicates with 10 technical replicates each.

**Supplementary Figure 3: Lower temperatures reduce HNEC colonization by CRS isolates of *S. aureus*.** Viable colony-forming units of *S. aureus* strains inoculated onto HNECs and incubated at 37°C (red) or 30°C (blue) for 6 hours. Two-way ANOVA \**P<*=0.05. Lines indicate matched biological replicates.

## Notes

### Competing Interest Statement

The authors have declared no competing interest.

