## Supplementary figures and images for "Temperature Influences Commensal-Pathogen Dynamics in a Nasal Epithelial Cell Co-culture Model"

### Supplemental figures

**Supplementary Figure 1.**

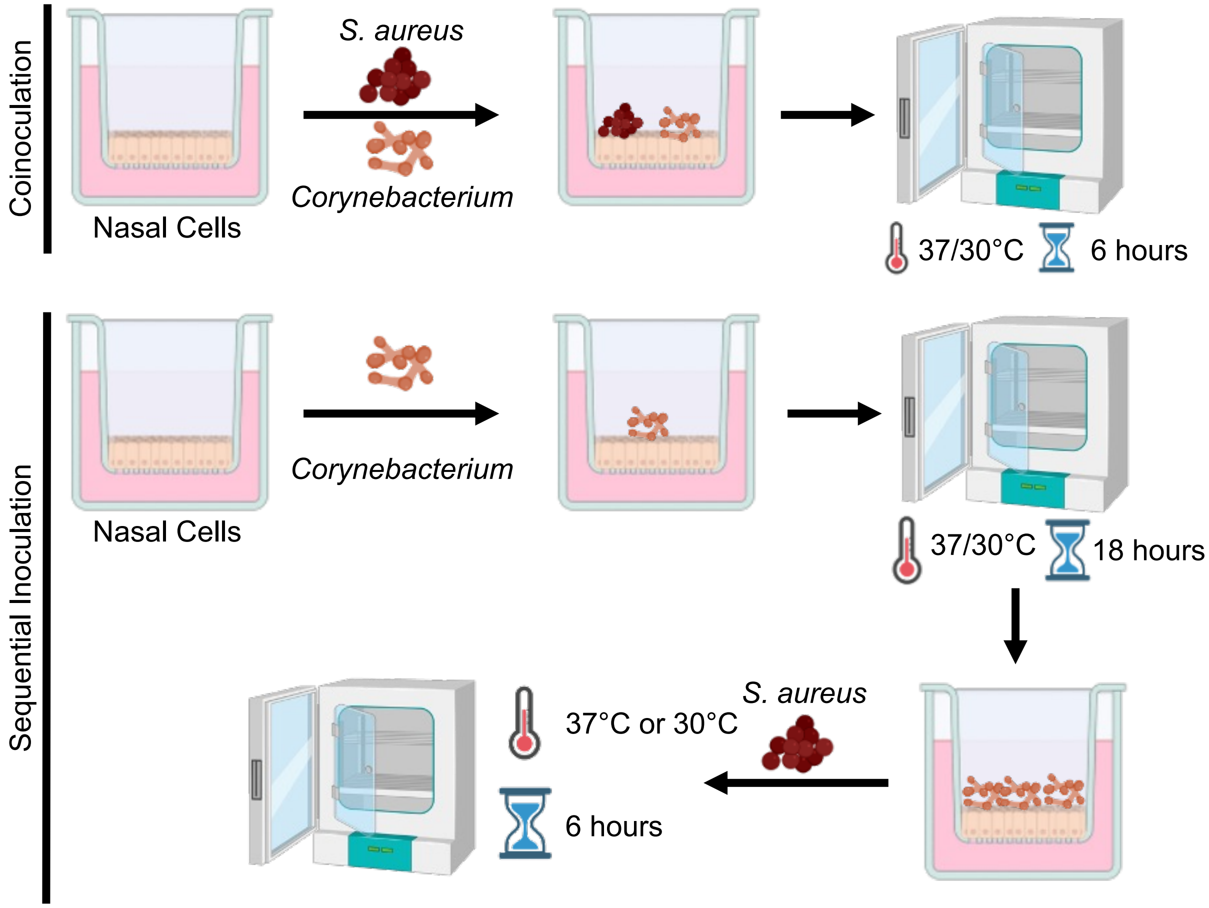

**Supplementary Figure 2.**

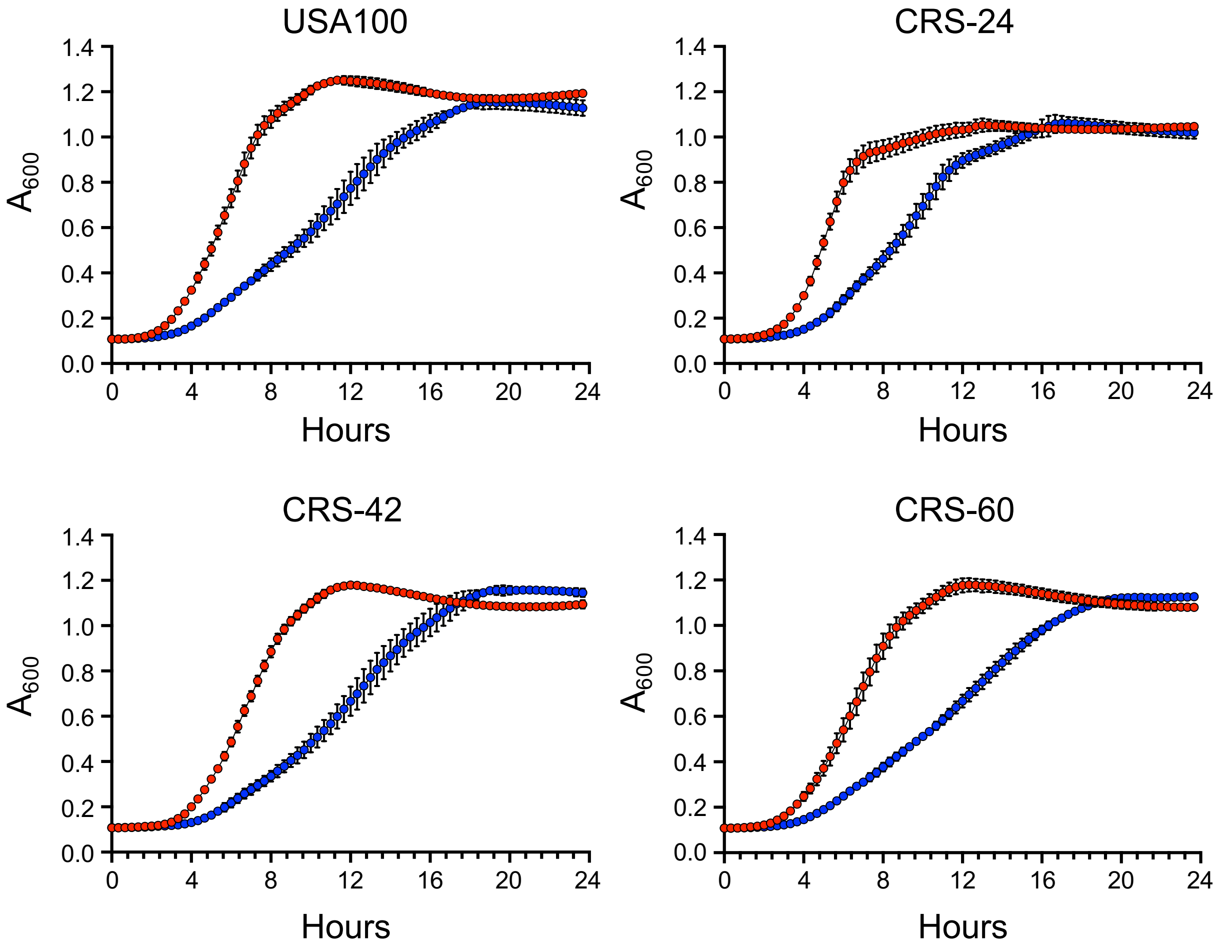

**Supplementary Figure 3.**

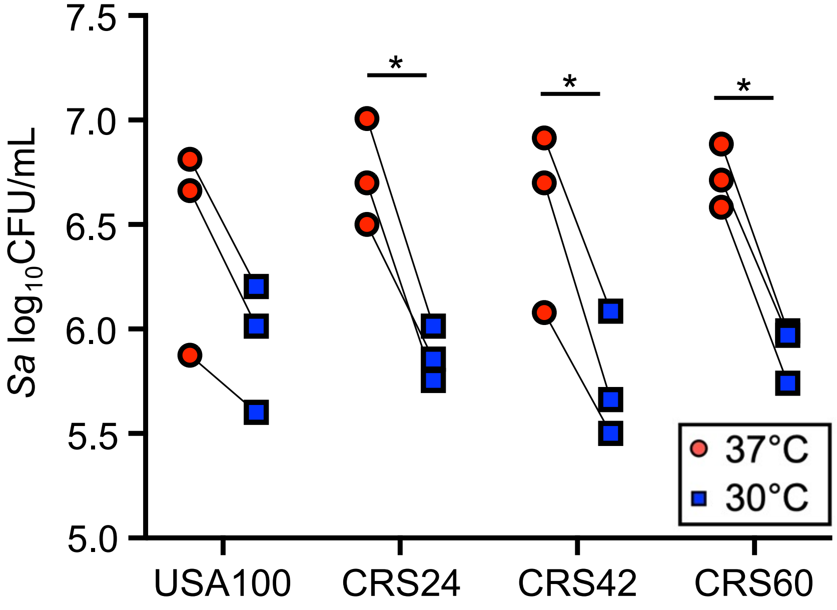
